# The Time Course of Ineffective Sham Blinding During 1mA tDCS

**DOI:** 10.1101/462424

**Authors:** Robert Greinacher, Larissa Buhôt, Lisa Möller, Gemma Learmonth

## Abstract

**Background:** Studies using transcranial direct current stimulation (tDCS) typically compare the effects of an *active* (10-30min) relative to a shorter *sham* (placebo) protocol. Both active and sham tDCS are presumed to be perceptually identical on the scalp, and thus represent an effective method of delivering double-blinded experimental designs. However, participants often show above-chance accuracy when asked which condition involved active/sham retrospectively.

**Objective/Hypothesis:** We aimed to assess the time course of sham-blinding during active and sham tDCS. We predicted that 1) Participants will be aware that the current is switched on for a longer duration in the active versus the sham protocol, 2) Active anodal tDCS will reduce reaction times more effectively than sham.

**Methods:** 32 adults were tested in a pre-registered, double-blinded, within-subjects design. A forced-choice reaction time task was undertaken before, during and after *active* (10min 1mA) and *sham* (20s 1mA) tDCS. The anode was placed over the left primary motor cortex (C3) to target the right hand, and the cathode on the right forehead. Two probe questions were asked every 30s: “*Is the stimulation on?* “and “*How sure are you?”.*

**Results:** Distinct periods of non-overlapping confidence intervals were identified between the active and sham conditions, totalling 5min (57.1% of the total difference in stimulation time). These began immediately after sham ramp-down and lasted until the active protocol had ended. Active tDCS had no effect on reaction times compared to sham (ΔRT active vs sham p>0.38 in all blocks).

**Conclusions:** We show a failure of placebo control during low-intensity tDCS.

## 1. Introduction

Transcranial direct current stimulation (tDCS) is a popular, non-invasive method of modulating cortical excitability. One of the main advantages of tDCS over other forms of neuromodulation, such as transcranial magnetic stimulation, is the ability to administer a placebo control protocol (*sham* tDCS) that is assumed to be perceptually indistinguishable from active stimulation on the scalp. In a typical sham protocol, the current is ramped-up gradually, then delivered for a short period at the same intensity as the active comparator (≤2 mins at 1-2mA), followed by a fade-out phase. This brief stimulation period seems not to induce any substantial neuromodulatory effects [1], but enables the initial cutaneous sensations associated with stimulation, including tingling, itching, burning and headache [2–4], to be presented in both the active and sham protocols.

Studies involving sham protocols therefore rely on participants being unable to differentiate the sensations experienced during this brief sham period from a prolonged period of active stimulation (10-20 mins). This is of vital importance to tDCS research, because a failure of sham-blinding could potentially encourage participants to modify their behaviour, even subconsciously, in response to knowing whether or not they are receiving an active protocol. This may be particularly compromising to a study where the hypotheses and outcome measures are communicated to participants beforehand, for example via the study information sheets [5]. The effectiveness of sham-blinding has been quantified in two broad ways in the prior literature: 1) by comparing the reported frequency and severity of cutaneous sensations during different protocols and 2) by asking participants to guess whether they had received active or sham tDCS (in the case of between-groups designs), or which of multiple sessions had involved active or sham (in within-group designs).

There is mixed evidence as to whether sham-blinding can indeed be successfully achieved during the application of standard tDCS parameters. Broadly, low-intensity (1-1.5mA) protocols can be blinded more effectively than high-intensity (2mA) tDCS [6]. However, the results are mixed, with some studies finding better-than-chance rates of guessing which condition had involved active/sham at both 2mA [6, 7] and 1-1.5mA [6, 8–11] current strengths, but others only around chance-level during high- [2, 12–15] and low- [2, 3, 16] intensity stimulation. However, these questions are typically probed on a retrospective basis, at the end of each session or experiment, and are therefore heavily reliant on the participants’ recollection of each event.

Indeed, there is scant evidence as to whether participants are able to dissociate active tDCS from sham *during* the course of an experiment. To assess this, Ambrus et al., (2012) probed the time course of cutaneous sensations at regular intervals during 10min of 1mA anodal stimulation, 10min of 1mA cathodal, and a 30s sham protocol. Participants were asked to rate the perceived strength of sensations felt on the scalp at 7 probe points, every 1.75min after tDCS onset (scale = “no sensation” to “extreme discomfort”). There was no difference in sensation strength between the three conditions at the first time point. The strength then reduced significantly in the sham condition from 2.25min onwards, in the anodal from 4 min and cathodal from 5.75min but there were no statistical differences between the three conditions. Specifically, Ambrus et al. (2012) argue that active and sham protocols are perceptually indistinct due to the persistence of sensations in the sham condition that endure after stimulation has ended. Although this study provides important information regarding the time course of cutaneous sensations during tDCS, we argue that the question posed does not adequately probe whether participants can tell that the stimulation is active at present. It is feasible that active and sham protocols may be similar in their subjective (dis)comfort ratings, yet participants still able to identify when the stimulation is active due to additional factors that are not probed by this question.

In this study, we aimed to assess the time course of sham-blinding by probing at 30s intervals whether participants can identify that 1mA tDCS is active or inactive, and the confidence of their decision. We chose to extend a well-powered, pre-registered reaction time experiment of Minarik et al. (2016), in which a small improvement in reaction time was elicited during 1mA of anodal tDCS to the primary motor cortex, relative to a cathodal protocol. Here we incorporated a 20s sham protocol instead of the cathodal condition. We hypothesised that 1) participants will be able to identify that a low-intensity current is switched on for a longer duration during the 10min active protocol, compared to the 20s sham, and 2) active anodal tDCS will reduce reaction times more effectively than sham.

## 2. Material and Methods

### 2.1 Pre-registration

The study protocol was pre-registered at https://osf.io/2zwhg/. The stimulus materials and full datasets are also available here.

### 2.2 Participants

An *a priori* sample size calculation, based on the effect size of d=0.45 observed in Minarik et al., (2016), identified that 32 participants were required for a one-tailed, repeated measures t-test, where power=0.8 and α=0.05 (Faul et al., 2009). Participants had a mean age of 24.5 years, range=19-38, and 27 were female. All were right handed, reported normal or corrected-to-normal vision, had no tDCS contraindications (as per Rossi et al., 2009) and had not received any form of electrical stimulation before. The study was approved by the University of Glasgow College of Science and Engineering ethics committee and written, informed consent was obtained from each participant.

### 2.3 tDCS

A direct current was applied using a battery-driven constant current stimulator (NeuroConn GmbH, Germany). Two protocols were administered in a double-blinded, counterbalanced, within-subjects design with ≥24 h between sessions: (i) ACTIVE ANODAL: 1mA tDCS for 10min, and (ii) SHAM: 1mA tDCS for 20s, both with additional 30s ramp-up and 30s ramp-down periods. In both protocols, the anode was placed vertically over the left primary motor cortex (C3 of the 10-20 International EEG system) and the cathode horizontally over the right forehead. Both carbon rubber electrodes measured 5×7cm and were encased in 0.9% NaCl saline-soaked sponges, held in place using rubber bands. The “study mode” of the NeuroConn stimulator was used to double-blind the protocols using 5-digit codes. During the sham protocol the device delivered a weak probe current of 110μA every 550ms to test the electrode impedance, which was relayed to the screen of the device. Electrode impedance was 6.2kΩ on average at the start of stimulation (range=4.1-10.7kΩ). Participants were given no information regarding the stimulation parameters (see information sheets provided at https://osf.io/2zwhg/).

### 2.4 Reaction time task and sham-blinding probes

The reaction time task was adapted from Minarik et al., (2016) and was performed on a Dell Precision T3400 PC and 19.5′′ Sun Microsystems CRT monitor with 1280×1024 pixel resolution and 100Hz refresh rate. The task in Block 1 (baseline) was identical to the procedure of Minarik et al., (2016): either a square or a diamond appeared in the centre of the screen for 100ms, followed by a fixation cross of variable duration between 1700-2100ms (Figure 1). Participants were instructed to press the left mouse button if they saw a diamond and the right mouse button for a square, as quickly and accurately as possible. The original stimuli of Minarik et al. were amended by affixing a diamond to the upper left corner and a square to the upper right throughout the experiment, as piloting highlighted that some participants erroneously switched responses mid-experiment. In Block 1, a total of 100 trials lasting 200s was presented. In Blocks 2-4, we inserted 2 probe questions into the reaction time task at 30s intervals, after every 10 trials, with each probe question presented on the screen for 4500ms. Question 1 involved a binary yes/no choice (“Is the stimulation on?”), and question 2 assessed the confidence of their decision (“How sure are you?”) via a visual analogue scale where 0=very unsure and 10=very sure. Questions were answered using a mouse click. The task was initiated in Block 2 at the same time as the 30s tDCS ramp-up began, with the first sham-blinding probe point occurring 30s after tDCS onset (i.e. at the point at which the stimulation reached 1mA in both the active and sham protocols). The task was then performed for 16 minutes continuously: during the 30s ramp-up, the 10 mins of online tDCS (active), the 30s ramp-down, then for 5 minutes offline after the active tDCS had ended. There were 220 trials during stimulation (Blocks 2&3) and a further 100 trials after stimulation had ended, providing a total of 32 probe points to assess the time course of sham-blinding.

**Figure 1.**
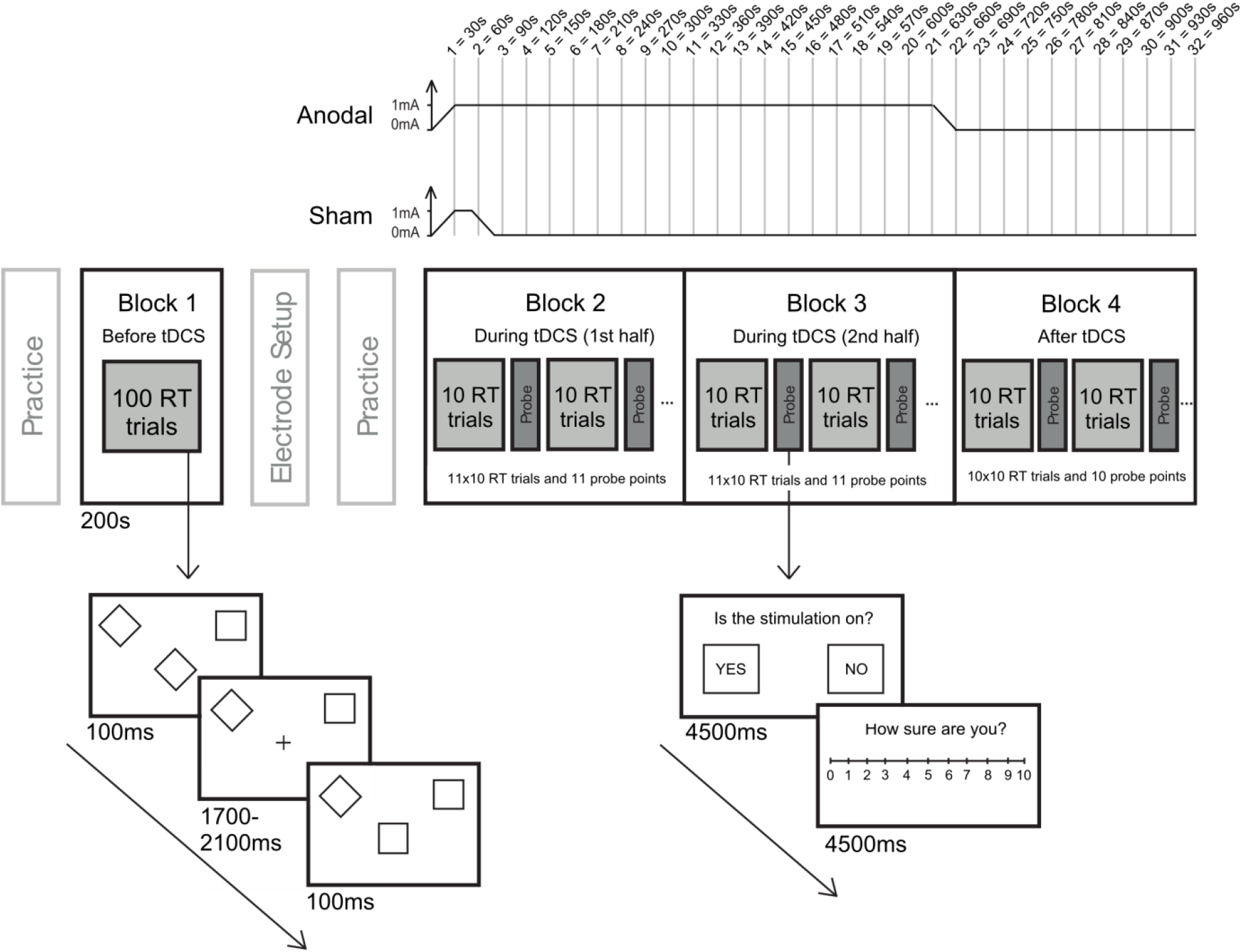
Illustration of the experimental design. Block 1 was performed as a baseline measure before tDCS onset. The stimulation ramp-up in both conditions began at the start of Block 2 and the active ramp-down finished at the end of Block 3. In Blocks 2-4 sham-blinding probe questions were inserted at 30s intervals to assess 1) whether participants could identify when the stimulation was switched on and 2) the confidence of their decision.

### 2.5 Procedure

Participants completed a short practice block (30 trials) of the reaction time task, followed by the baseline Block 1. The electrodes were then applied to the scalp. The tDCS device was programmed using a pre-allocated 5-digit code and the resistance was checked and lowered if necessary. A short practice of the reaction time task including the sham-blinding probes was performed. Block 2 of the behavioural task was initiated at the same time that the stimulator began its ramp-up to 1mA. Electrode resistance was recorded when the tDCS device reached 1mA. Blocks 2-4 were then completed in a single, continuous 16min period. The stimulation ramp-down coincided with the end of Block 3 in the active condition. Block 4 was performed after active tDCS. After the electrodes were removed, participants rated whether they had experienced headache, tingling, itching, burning or pain during the session (1=Not at all, 5=Very strongly). At the end of the second session they were informed about the two protocols and asked to guess which of the two sessions had involved sham, and the confidence of their guess (scale=1-10). In contrast to the between-groups design of Minarik et al. (2016), here participants attended on two different days, a minimum of 24 hours apart, in a within-subjects design. Finally, after all 32 participants had been tested, the researchers were un-blinded to the stimulation sessions and the data sorted into active and sham sessions.

## 3. Results

### 3.1 Effectiveness of sham-blinding

We had pre-registered a plan to fit binomial logistic curves for each individual and calculate the time point within the experiment at which a 50/50 yes/no guess rate was reached. Unfortunately, the data was found to be unconducive to this analysis due to participants switching between yes and no responses more frequently than anticipated. It should therefore be noted that these results are derived from a post-hoc alteration to the analysis plan. The outcome measures derived from the two probe questions (*Is the stimulation on?* and *How sure are you?*) were combined to create a weighted score. A “yes” response to the first question was assigned a value of +1 and “no” a value of −1. This was then multiplied by the confidence rating (0-10), so that a value of +10 indicated high confidence that the stimulation was switched on, and −10 high confidence that it was switched off. We then bootstrapped 95% confidence intervals for each of the 32 probe points, separately for the active and sham conditions, using 5000 permutations of the data (Figure 2).

**Figure 2.**
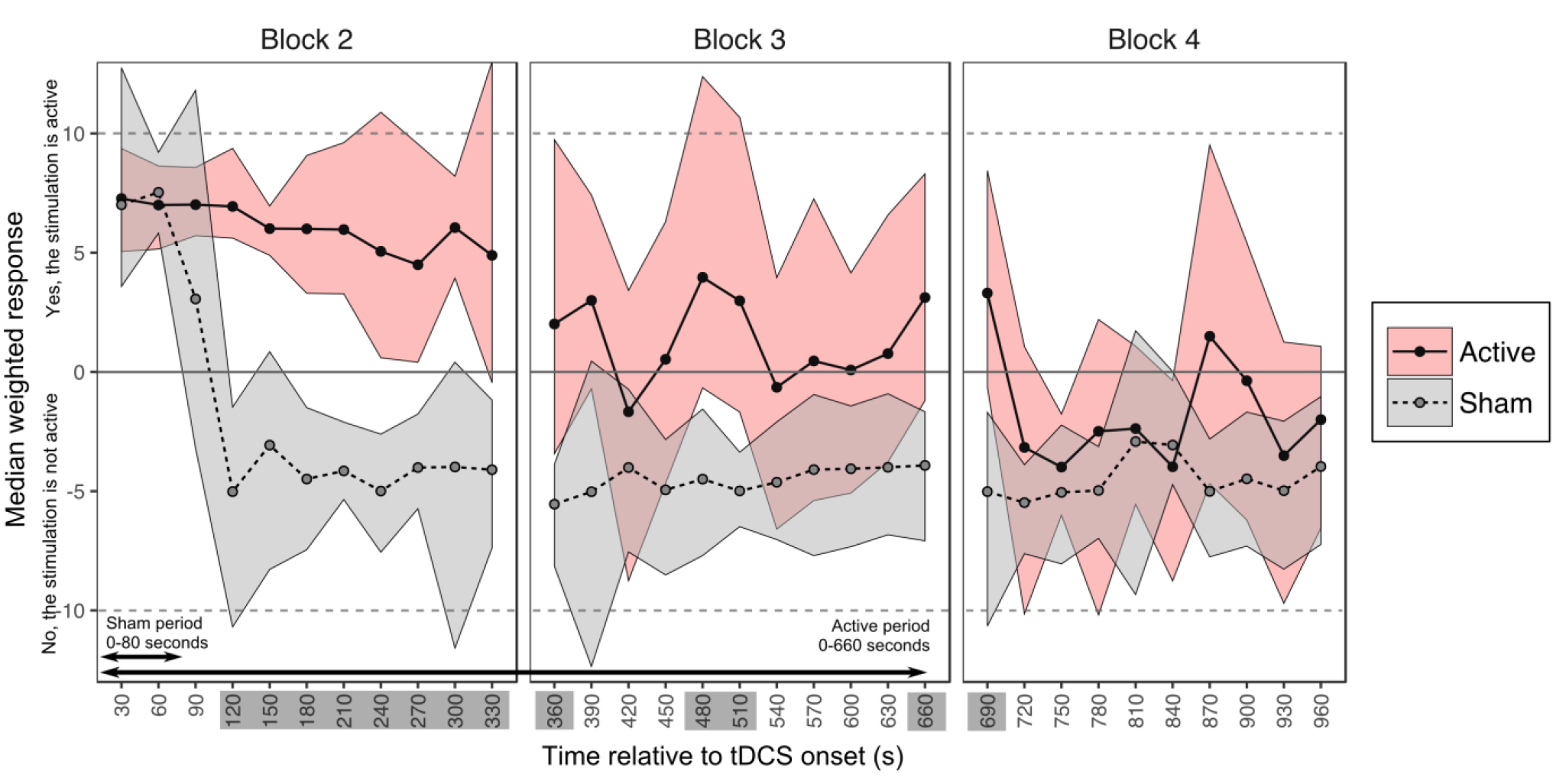
Sham-blinding results. The median weighted responses are shown for active and sham, with bootstrapped 95% confidence intervals. Time points with distinct (i.e. non-overlapping) ratings for active and sham are highlighted.

We identified overlapping confidence intervals between the active and sham protocols throughout the first 90 seconds of stimulation, where participants were confident that the tDCS was switched on in both conditions. This period included the 30s ramp-up in both protocols, followed by the 20s 1mA sham tDCS and the 30s sham ramp-down. In the active condition, confidence intervals remained above zero until 5 min (300s) post-tDCS onset. Group-level responses became more varied from this point onwards, with wider confidence intervals that overlapped zero, although the median value generally remained above zero until the active stimulation had ended. Conversely, in the sham condition, participants were confident that no stimulation was being delivered from 2 min (120s) post-tDCS onset and this lasted broadly until the end of the experiment. Comparing active and sham directly, the confidence intervals were distinct for a total of 5 min and this represented 51.7% of the total difference in stimulation time between conditions (see highlighted time-points on the x-axis of Figure 2). The first non-overlapping period started at 120s post-tDCS onset and lasted until 360s, a second period was identified between 480-510s and a third between 660-690s (the active protocol had fully ramped-down by 660s). Finally, the weighted scores were no different between active and sham during Block 4, when the stimulation had ceased in both protocols, from 12min (720s) until the end of the experiment.

### 3.2 Reaction times

Participants were highly accurate in the behavioural task throughout the experiment (mean accuracy=95.6%, bootstrapped 95% CI=[95.08, 96.11%]). As per the pre-registered analysis plan, the median response time for correct trials was calculated for each participant per block (Figure 3A). A TOST procedure indicated that baseline reaction times were equivalent in both conditions (i.e. the observed effect size of d_z_= −0.02 was within the equivalent bounds of a moderate effect size of d_z_= −0.4 and 0.4; t(31)=2.15, p=0.02 [20]). The baseline RTs (Block 1) were then subtracted from each subsequent block to create a ΔRT value for each of Blocks 2-4 (Figure 3B). A series of three one-tailed, repeated-measures t-tests were performed to compare the ΔRT in the active and the sham protocols, separately per block. There were no differences in RT shifts from baseline between the active and sham conditions in any of the 3 blocks (ΔRT Block 2: t(31)=0.26, p=0.4, d=0.042; Block 3: t(31)=0.16, p=0.44, d=0.028; Block 4: t(31)=0.29, p=0.39, d=0.047). As an exploratory follow-up, we assessed whether there were any transient differences between anodal and sham tDCS that might have been obscured by collapsing the reaction time data into 5-min blocks. The median reaction times were bootstrapped for each sub-block of 10 trials in Blocks 2-4, but no non-overlapping periods were observed between conditions (Figure 3C).

**Figure 3.**
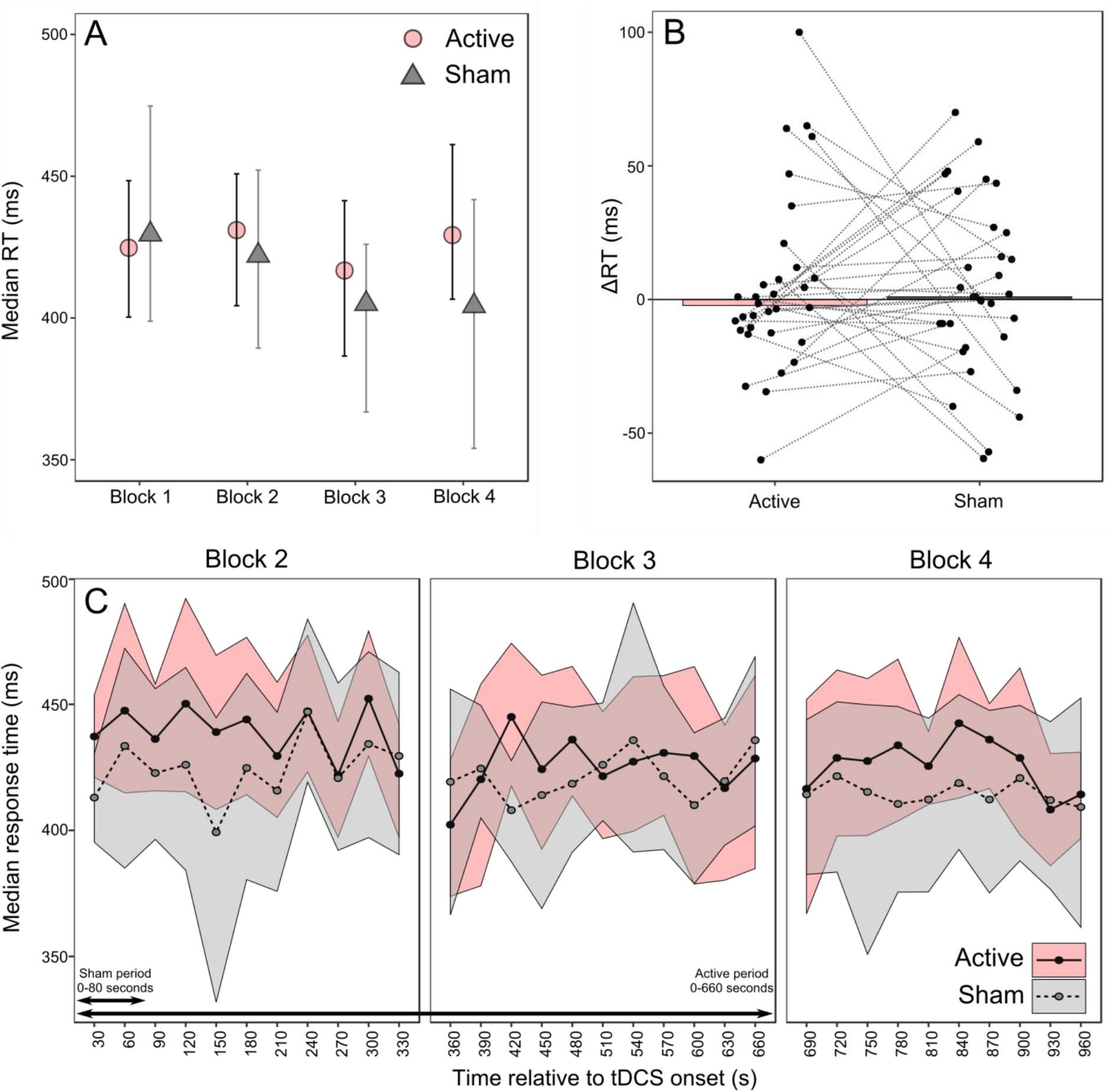
Reaction time results. A) Median group-level RTs per block with bootstrapped 95% CIs. B) Median change in RT from baseline to Block 3 with individual participant medians shown. C) Median RTs for each sub-block of 10 trials with bootstrapped 95% CIs.

### 3.3 Side-effects and sham-blinding questionnaires

A series of Wilcoxon signed rank tests identified a significant difference in the retrospective rating of itchiness between the active and sham conditions (Z= −3.13, p=0.002). There were no differences for ratings of headache (Z= −0.18, p=0.86), tingling (Z= −1.29, p=0.197), burning (Z= −1.25, p=0.21) or pain (Z= −0.979, p=0.33). Participants were above chance (25/32=78.1%) in their ability to guess which of the two sessions had involved sham at the end of the experiment.

## 4. Discussion

We present here two main findings. Firstly, participants were confident that active stimulation was being delivered for a longer duration during 10mins of active anodal tDCS compared to a 20s sham protocol. Secondly, 10 min of 1mA anodal tDCS to the primary motor cortex failed to reduce reaction times relative to sham. Our results highlight a failure of sham-blinding during low-intensity tDCS, adding to previous similar results at a higher current [6] and adding important information regarding the time-course of failed sham-blinding. Specifically, we show that the cessation of stimulation in the sham protocol resulted in an immediate and sharp increase in the number of participants who were confident that the stimulator had been deactivated, which lasted until the end of the experiment. It is important to note that participants were given no prior information about the two stimulation conditions and the experimenters were also blinded to the counterbalancing across conditions. These results contradict prior claims that active and sham protocols are perceptually indistinct due to a persistence of sensations after the ramp-down has ended in sham protocols [16]. In the active protocol, we found a maintained confidence that the stimulation was still switched on until 5 mins into the stimulation period, and periods of difference relative to sham that lasted broadly until the stimulation had ended in the active condition. We did, however, observe a gradual reduction in group-level confidence over the course of the 10 min anodal stimulation, supporting the general concept of scalp desensitisation during sustained active tDCS. However, we argue that placebo controls must remain consistently blinded over the whole time-course of application to be deemed effective, particularly if participants are made aware of protocol differences beforehand [5, 21].

Our findings provide new and important information regarding the time course of sham-blinding that could only be revealed by probing this issue throughout the experiment, rather than retrospectively. Studies that probe whether sham-blinding was achieved on a retrospective basis could potentially introduce a number of confounding variables that may influence the accuracy of these reports. Firstly, participants may have a poor recollection of what they experienced during each session, particularly if there were multiple sessions with long periods between testing days. Secondly, the probe questions may not allow for nuanced responses relating to the certainty of these decisions and what factors might have led to them, nor the duration of any side-effects experienced. For instance, we often anecdotally find that participants report the tightness of the rubber bands to be the most uncomfortable aspect of receiving tDCS. It is possible that participants could assign a disproportionate weighting to such factors in their reports, which are unrelated to stimulation *per se*, and would likely be identical across tDCS conditions. Such factors could account for the mixed reports in the prior literature [2,6–10, 12–15]. In going some way to address this issue, Ambrus et al. (2012) found no difference in the time-course of *discomfort* ratings between active and sham tDCS, which they interpreted as successful sham-blinding. However, here we find prolonged differences when participants are asked specifically whether or not they believed the stimulator to be active.

It is important, then, to consider possible methods of delivering more effective sham blinding during electrical stimulation studies. Topical anaesthetics can reduce or eliminate skin sensations associated with tDCS [22, 23], although might mask more serious side-effects such as skin burning [24]. However this appears to be low-risk using standard stimulation parameters [25]. A second option is to compare anodal stimulation against a cathodal paradigm, without including a sham, which should give rise to observable differences in outcome measures. This approach is often precluded e.g. in certain clinical groups such as stroke, where it would be counterintuitive to deliver a protocol which further impedes behaviour. A third option is to compare the outcomes of two different electrode montages, one targeting the cortical area of interest and the second a location uninvolved in that behaviour (“*off-target active stimulation*” [21]). It is also possible to reduce priming effects caused by participant expectations by using *de facto* masking, where participants are erroneously informed that the stimulation is active in all sessions [6]. However, it is unclear whether this form of masking could successfully counteract the sensory differences observed between protocols as described here. In fact, our participants were informed that “weak electric current will be applied over your scalp for several minutes” but they remained able to tell when the stimulation ended during the sham protocol. We suggest that the appropriate control should be carefully tailored based on the specific aims and participant population of each individual study. We also recommend that researchers ensure that the aims, hypotheses and study design are concealed in any information sheets provided to participants to minimise demand characteristics associated with identifying the active stimulation condition.

As a secondary finding, we also failed to find a difference in reaction times during anodal tDCS relative to sham, adding to previous null results relating to stimulation of the primary motor cortex [26–30]. Since Minarik et al. (2016) did not include a sham protocol, it was not possible to determine whether the anodal tDCS resulted in improved reaction times, or whether the cathodal protocol had actively inhibited motor learning throughout the task. We found that our sham protocol elicited a similar change in reaction time to their cathodal condition, but here we failed to find a facilitation of reaction times in response to anodal tDCS. However, we should note that we modified both the task and experimental design in the present study. After baseline testing, Minarik et al. (2016) delivered 4 min of tDCS offline, prior to 4 min online tDCS, during which reaction times were re-assessed. Our active protocol was delivered entirely online, and there is current debate regarding whether anodal stimulation is more effective in modulating motor cortex excitability when applied before, rather than during, behavioural tasks [31].

It could be argued that by directly asking participants whether the stimulation is active, we may have drawn attention to this issue more than in a typical experiment. However, we reason that the effectiveness of placebo control should not be reliant upon participants being able to distract themselves from thinking about the conditions that are being administered. Secondly, it is empirically unknown what participants *do* think about during stimulation experiments, and this factor cannot be easily tested nor controlled. Although participants are instructed to focus on a task during online tDCS delivery, they may be simultaneously thinking about the sensations on their scalp. We would then also expect to find higher rates of failed sham-blinding in offline experiments, when participants are instructed to rest during stimulation with no task to distract from these sensations. A second consideration is that our participants experienced both stimulation conditions and were able to compare sensations across days. However, within-subjects and crossover designs are common in electrical stimulation studies, particularly in clinical trial designs, and thus achieving sham-blinding in these designs remains a pertinent issue. We recommend that further, basic methodological studies be conducted to develop improved sham protocols that can be adequately blinded from active electrical stimulation.

## 5. Conclusions

We conclude that the standard method of sham-blinding reported here may be ineffective, and could allow participant expectations to influence the study outcomes. These results highlight a need to develop more effective methods of delivering blinded placebo control protocols during electrical stimulation experiments.

## Acknowledgements

This work was supported by a Sir Henry Wellcome Postdoctoral Fellowship awarded to GL (209209/Z/17/Z). We wish to thank Dr Monika Harvey for use of the tDCS equipment.

